# Hydrogen sulfide-mediated vasodilation requires heme oxygenase-derived carbon monoxide

**DOI:** 10.64898/2026.08.11.744278

**Authors:** Jacob R Anderson, Crystal X Nguyen, Laura V Gonzalez Bosc, Jay S Naik

## Abstract

**Background:** Hydrogen sulfide (H_2_S) is an important endothelial-derived vasodilator, but the signaling mechanism remains incompletely understood. We previously demonstrated that H_2_S-mediated vasodilation requires transient receptor potential vanilloid type 4 (TRPV4) channels. Because H_2_S has been reported to enhance heme oxygenase (HO) activity and HO-derived carbon monoxide (CO) regulates endothelial signaling, we hypothesized that H_2_S-mediated vasodilation requires HO-2-derived CO.

**Methods:** Pressure myography was performed in isolated rat mesenteric arteries to determine the contribution of HO, TRPV4, eBK, and SK/IK channels to H_2_S-mediated vasodilation. HO-2 sulfhydration was assessed using a maleimide assay, and spatial association among HO-2 and TRPV4 was examined using proximity ligation assays in human aortic endothelial cells.

**Results:** H_2_S Selicited concentration-dependent vasodilation that was abolished by HO inhibition. Repletion of CO restored H_2_S-mediated vasodilation in the presence of HO inhibition. CO-mediated vasodilation was abolished by TRPV4 and SK/IK inhibition but was unaffected by eBK inhibition. H_2_S increased HO-2 sulfhydration and enhanced HO activity. In endothelial cells, HO-2 and TRPV4 exhibited close spatial association.

**Conclusions:** These findings support a model in which H_2_S stimulates HO-2-derived CO production, leading to TRPV4-dependent endothelial signaling, SK/IK activation, and vasodilation. Together, the data support the existence of an endothelial HO-2/TRPV4/SK/IK signaling domain that contributes to H_2_S-mediated vascular reactivity.

## 1 Introduction

The endothelium plays a critical role in regulating vascular tone by integrating mechanical and chemical stimuli and releasing vasoactive signaling molecules. Among these signaling mediators, the gaseous transmitters nitric oxide (NO) (Joyner & Dietz, 1997), carbon monoxide (Caudill et al., 1998; Naik et al., 2003; Naik & Walker, 2001, 2003), and hydrogen sulfide (H_2_S)(Cirino et al., 2023; Jackson-Weaver et al., 2013; Mendiola et al., 2022; Morales-Loredo et al., 2019a; Naik et al., 2016) have emerged as important regulators of vascular homeostasis. While NO is widely recognized as an important mediator of endothelium-dependent vasodilation in many vascular beds, increasing evidence supports physiological roles for both CO and H_2_S in regulating blood pressure and organ blood flow (Choi & Kim, 2021; Cirino et al., 2023; Mendiola et al., 2021; G. Yang et al., 2008).

H_2_S is produced within the vasculature by the enzymes cystathionine-lyase (CSE) and 3-mercaptopyruvate sulfurtransferase (Cirino et al., 2023). Previous work has shown that diminished H_2_S production is associated with the development of hypertension in humans and animal models (Jackson-Weaver et al., 2011; Morales-Loredo et al., 2019a; Sun et al., 2007; C. Wang et al., 2014). For example, hypertension develops with age in mice genetically deficient in CSE (G. Yang et al., 2008). Rajpal et al. found that the frequency of the CTH/CSE 1364 G > T allele was significantly higher in patients with cardiovascular disease than in the control group, suggesting that this single-nucleotide polymorphism may be a potential genetic risk factor (Rajpal et al., 2018). We have shown that endothelial cells (EC) are the source of endogenous vascular H_2_S production and are required for H_2_S-mediated vasodilation (Jackson-Weaver et al., 2013; Naik et al., 2016). Previous work from our laboratory demonstrated that H_2_S-mediated vasodilation in resistance-sized mesenteric arteries requires endothelial TRPV4 activation (Naik et al., 2016) and calcium-activated potassium channels (Jackson-Weaver et al., 2013; Liang et al., 2012; Mendiola et al., 2022; Naik et al., 2016). However, the signal transduction pathways underlying H_2_S-mediated vasodilation remain incompletely defined.

Heme oxygenase (HO) catalyzes the conversion of heme to biliverdin, releasing CO as a byproduct. HO exists as three isoforms: inducible (HO-1), constitutive (HO-2), and a third not yet fully characterized form (HO-3) (Wilks, 2002). We have previously shown that both HO-1 and HO-2 are expressed in ECs of the aorta and mesenteric arterioles in rats (Naik et al., 2003). CO can activate signaling pathways that promote vasorelaxation and suppress vascular contractility (Leffler et al., 2011). Previous studies have demonstrated that HO-derived CO modulates vascular tone in both physiological and pathophysiological settings (Caudill et al., 1998; Naik et al., 2003; Naik & Walker, 2003). Furthermore, endothelial HO-derived CO has been implicated in regulating potassium channel activity and endothelial signaling pathways associated with vasodilation (Bolognesi et al., 2007; Dong et al., 2007; Leffler et al., 2011; Riddle & Walker, 2012; Xi et al., 2004).

Emerging evidence suggests cross-talk between H_2_S and CO signaling pathways. H_2_S has been reported to enhance HO activity by interacting with ferric verdoheme intermediates (Matsui et al., 2018), thereby increasing CO production and binding to multiple heme proteins (Boubeta et al., 2020). In addition, CO signaling has been linked to TRPV4-mediated Ca^2+^ influx (F. Yang et al., 2016), suggesting a potential mechanistic link among H_2_S, HO, and TRPV4. Taken together, these findings suggest that HO-derived CO may serve as an intermediary signaling molecule that couples H_2_S to TRPV4-dependent endothelial hyperpolarization and vasodilation. However, the contribution of HO-derived CO to H_2_S-mediated vasodilatory signaling has not been directly examined. Therefore, the present study tested the hypothesis that H_2_S-mediated vasodilation requires HO-derived CO-mediated TRPV4 activation.

## 2 Methods

Male Sprague–Dawley rats (Envigo, 250–300 g, 9–10 weeks old) were used for all experiments. Rats were euthanized with pentobarbital sodium (200 mg/kg I.P.), and mesenteric arteries were collected for experiments. The Institutional Animal Care and Use Committee of the University of New Mexico School of Medicine reviewed and approved all animal protocols. All protocols conformed to National Institutes of Health guidelines for animal use.

### Vasodilation Studies

Third- or fourth-order (small, lumen diameter < 200 µm) mesenteric artery segments were isolated in chilled HEPES-buffered physiological saline (HPSS; in mM: 130 NaCl, 4 KCl, 1.2 MgSO_4_, 4 NaHCO_3_, 1.8 CaCl_2_, 10 HEPES, 1.18 KH_2_PO_4_, and 6 glucose; pH adjusted to 7.4 with NaOH) and cleaned of adipose tissue. Arteries were then cannulated and pressurized to 60 mmHg in a single-vessel chamber (CH-1, Living Systems), equilibrated in HPSS for 15 min, and superfused at 5 mL/min at 37 °C. Mesenteric arteries from naive rats exhibit little myogenic tone (Jackson-Weaver et al., 2011). Therefore, following equilibration, arteries were preconstricted to 30-50% using U-46619 0.1 – 0.2 μM), and vasodilation was measured during the cumulative addition of the H_2_S donor NaHS (1, 10, and 30 M), CROM-3 (CO donor; 0.1, 1, and 10 nM), the TRPV4 agonist GSK1016790A (1 and 10 nM), or Acetylcholine (1 μM). Subsets of experiments were performed in the presence or absence of the following inhibitors, a heme oxygenase inhibitor, chromium mesoporphyrin (1 μM CrMP), the TRPV4 antagonist GSK2193874 (300 nM), the large conductance (BK) Ca^2+^-activated K^+^ channel inhibitor, iberiotoxin (100 nM, IBTx), or the small (SK) and intermediate conductance (IK) Ca^2+^-activated K^+^ channel inhibitors, apamin (100 nM) and TRAM-34 (1 μM), respectively. Arterial inner diameter was recorded using edge-detection software (IonOptix).

### Endothelial cell Ca^2+^ measurements

Isolated, cannulated, and pressurized mesenteric arteries (60 mmHg) were loaded intraluminally with fluo-4 AM (5 μmol/l, Invitrogen) containing 0.25% pluronic acid in HEPES buffer (PSS; in mmol/l: 129.8 NaCl, 5.4 KCl, 0.83 MgSO_4_, 0.43 NaH_2_PO_4_, 19 NaHCO_3_, 1.8 CaCl_2_, and 5.5 glucose) for 15 min at 28°C. After equilibration, endothelial cell Ca^2+^ events were assessed by exciting the fluorophores with a solid-state 488 nm laser (50% laser intensity), and emitted light >500 nm was collected using an Olympus IX83 inverted microscope with a Martzhauzer ultra-fast motorized XY stage with a 60X silicon-immersion lens and a spinning-disk confocal scanning unit (Yokogawa CSU-W1). Time-series images were acquired before and after H_2_S or CORM-3 in the presence or absence of CrMP or GSK219 at approximately 36 frames/s (500 images over 14 s). Ca^2+^ event analysis was performed using the ImageJ plugin XY-Spark.

### Proximity Ligation Assay

Protein-protein localization in human aortic ECs was determined using Duolink in situ proximity ligation assay (PLA) according to the manufacturer’s instructions. Human aortic ECs (HAoECs; Cell Applications 304-05a) were grown to confluence on an 18-chamber ibidi slide coated with fibronectin (1 μg/mL, Sigma FC010). HAoECs were fixed using 4% PFA and incubated with Duolink blocking buffer for 30 min at 37 °C before incubating overnight with mouse anti-HO-2 (1:100, Santa Cruz Biotechnology [sc-17786], RRID: AB_62772) and rabbit anti-TRPV4 (1:100, Proteintech [84091-4-RR], RRID: AB_3671657). HAoECs were then incubated with the appropriate PLA probes (1:5) for 1 h at 37 °C. Samples were mounted with Prolong Gold anti-fade with DAPI (Invitrogen P36935), and images of the PLA interactions were acquired using a confocal microscope (Zeiss LMS800 AxioObserver inverted microscope with X-Y motorized stage). Negative controls were performed by omitting the primary antibody (not shown).

### Sulfhydration of HO-2

Sulfhydration of HO-2 was measured using a previously described method (Sen et al., 2012). Briefly, rat testes were harvested and homogenized using lysis buffer containing Halt Protease inhibitor and EDTA (Invitrogen). Tissue lysates were treated with vehicle or 10 μM NaHS for 30 min at 37°C. Cells were lysed and homogenized, and HO-2 was immunoprecipitated using A/G Beads (Invitrogen). 2% of the input from each sample was reserved to determine total HO-2 protein using a standard Western blot. Samples were incubated with Alexa Fluor 680-conjugated C2 maleimide (2 μM) for 2 h at 4°C, followed by incubation with or without DTT (1 mM) for 1 h at 4°C. Samples were subjected to gel electrophoresis, transferred to a nitrocellulose membrane, and scanned with the LI-COR Odyssey system. Images were quantified using ImageJ. The fluorescence level in each sample was normalized to the total HO-2 protein level.

### In vivo HO Activity

Under isoflurane anesthesia, male Sprague-Dawley rats were infused intravenously through the jugular vein with either saline or Na₂S at a dose of 0.6 mg/kg/min for 20 min. A total infusion volume of 100 µL was delivered for each animal. Following infusion, blood was collected by cardiac puncture and allowed to coagulate at room temperature for 30 min. Samples were then centrifuged at 3,000 rpm for 15 min to isolate serum. Total serum bilirubin concentration was measured using a colorimetric bilirubin assay kit (Sigma-Aldrich, MAK126) according to the manufacturer’s instructions. Data were collected from five independent experiments for each treatment group.

### Statistical Analysis

Data are presented as means ± SD and were analyzed using two-way ANOVA or Student’s t-test, as appropriate (GraphPad Prism). P < 0.05 was considered statistically significant for all analyses.

### Reagents

U46619, GSK1016790A, and Iberiotoxin were purchased from Cayman Chemicals (Ann Arbor, MI). TRAM-34 and Apamin were purchased from MedChemExpress (Monmouth Junction, NJ). Chromium mesoporphyrin was purchased from Frontier Specialty Chemicals (Logan, UT). CORM-3 was purchased from Torcis BioSciences (Minneapolis, MN). All other reagents were purchased from Sigma-Aldrich (St. Louis, MO).

## 3 Results

### H_2_S-mediated vasodilation requires heme oxygenase

To determine whether HO signaling contributes to H_2_S-mediated vasodilation, isolated pressurized rat mesenteric arteries (150–200 μm) were exposed to increasing concentrations of the H_2_S donor NaHS in the presence or absence of the HO inhibitor, CrMP. NaHS elicited concentration-dependent vasodilation (Figure 1A) and increased endothelial calcium events (Figure 1B) under control conditions. Inhibition of HO significantly attenuated H_2_S-mediated vasodilation and endothelial Ca^2+^ events across all concentrations tested, indicating that HO activity is required for H_2_S-mediated vasodilation. Repletion of CO with a sub-dilator concentration of the CO donor CORM-3 restored NaHS-mediated vasodilation in the presence of CrMP, indicating that CO can rescue the loss of HO activity. In addition, these findings further demonstrate the specificity of CrMP and suggest that biliverdin and its downstream metabolites do not contribute to the observed response. Acetylcholine-induced vasodilation was not significantly altered by either CrMP or CORM-3 treatment (Supplemental Figure 1). Together, these findings support a model in which H₂S-mediated vasodilation requires HO activity and downstream CO-dependent signaling.

**Figure 1.**
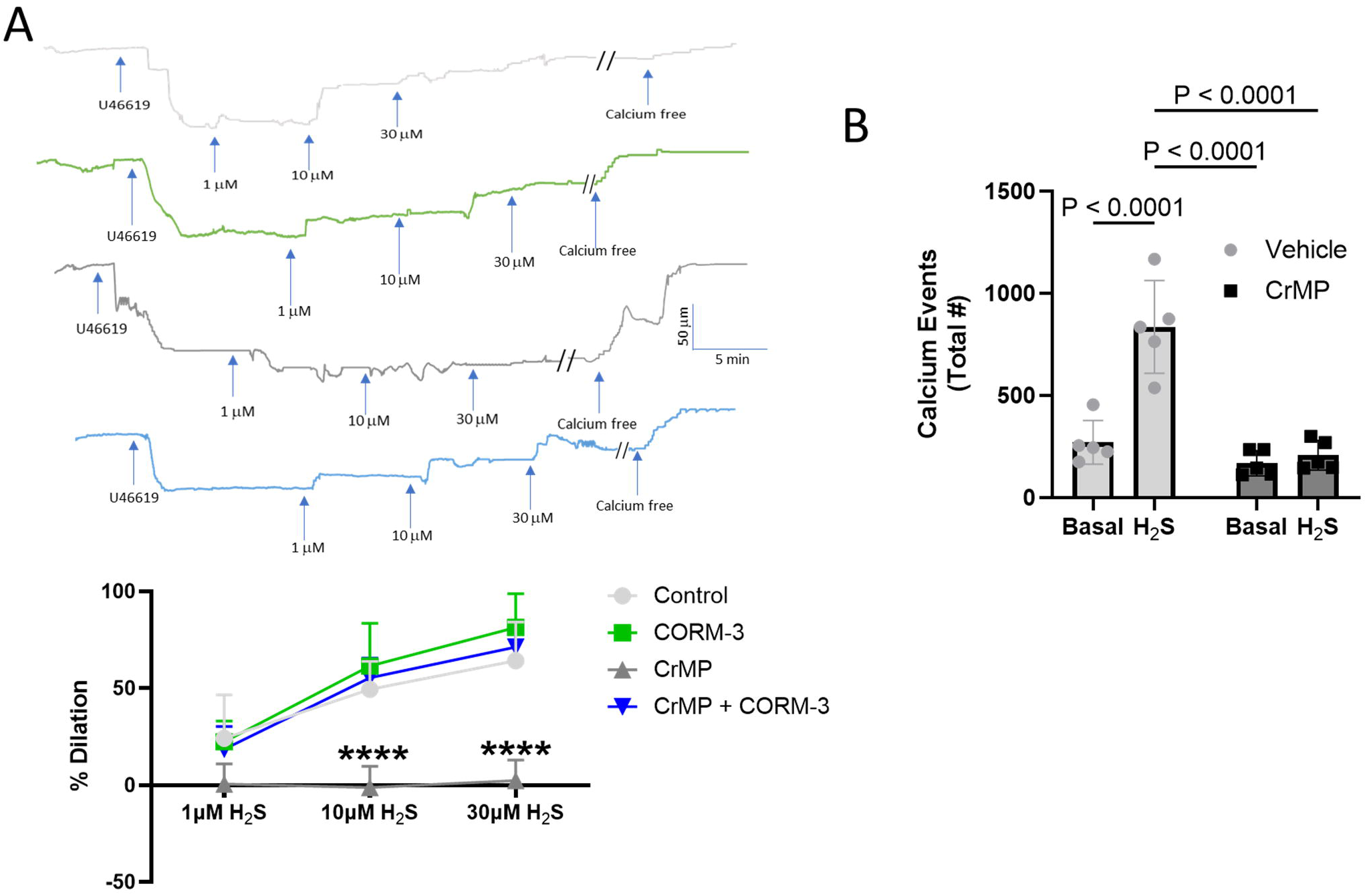
HO-2 inhibition attenuates H₂S-induced vasodilation and endothelial Ca²⁺ signaling in pressurized rat mesenteric arteries (150 – 200 μm). (**A**) Representative traces and summary data. Concentration-dependent vasodilation to NaHS (1, 10, and 30 µM) was assessed in isolated, pressurized mesenteric arteries (150–200 µm) under control conditions and in the presence of the HO-2 inhibitor chromium mesoporphyrin IX (CrMP; 1 µM) or the CO donor CORM-3 (10 pM). (**B**) Endothelial cells of pressurized mesenteric arteries were loaded with the fluorescent Ca²⁺ indicator Fluo-4 AM (5 µM), and endothelial Ca²⁺ events were measured under baseline conditions and following exposure to NaHS (10 µM), in the presence or absence of CrMP (1 µM). Data are presented as mean ± SD (n = 5/group) and were analyzed by two-way ANOVA. ****P < 0.0001 for CrMP versus all other conditions.

### H_2_S increases HO-2 sulfhydration and enhances HO activity

A maleimide sulfhydration assay was performed in rat testis lysates treated with NaHS or vehicle. Fluorescently labeled maleimide binds to sulfhydryl groups. Treatment with DTT reduces only sulfhydrated cysteines, resulting in a decrease in the fluorescent signal in proteins containing sulfhydrated residues. Exposure to NaHS increased HO-2 sulfhydration compared with vehicle-treated samples (Figure 2A). We have previously demonstrated that intravenous administration of the H_2_S donor Na_2_S decreased mean arterial pressure in conscious rats (Morales-Loredo et al., 2019b). Intravenous administration of Na_2_S increased serum bilirubin levels (Figure 2B). These findings are consistent with the hypothesis that H₂S-dependent sulfhydration of HO-2 contributes to increased HO activity.

**Figure 2.**
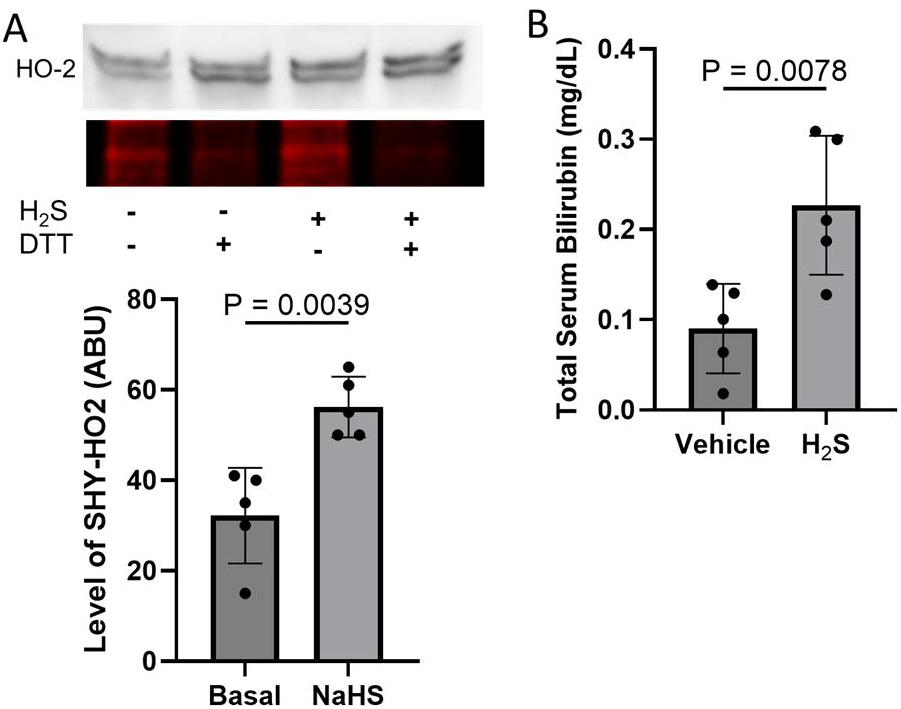
H_2_S sulfhydrates HO-2 and increase total serum bilirubin. (**A**) Sulfhydration detected by maleimide assay. Rat testis lysates were treated with vehicle or 10 μM NaHS for 30 min at 37°C. Lysates were immunoprecipitated overnight at 4°C and labeled with maleimide. Samples were divided into two conditions: either reduced with DTT (+DTT) or vehicle (−DTT). Maleimide fluorescence was normalized to total HO-2 protein determined by Western blot. (**B**) Intravenous infusion of Na_2_S (0.6 mg/kg.min) for 20 min increased total serum bilirubin. Data are presented as mean ± SD (n = 5/group) and analyzed by unpaired Student’s t-Test.

### HO/CO system is upstream of TRPV4 activation

We previously demonstrated that H_2_S can sulfhydrate TRPV4 channels, increase TRPV4-mediated Ca^2+^ events, and elicit TRPV4-dependent vasodilation (Naik et al., 2016). In the present study, H_2_S sulhydrated HO and increased HO activity. To determine whether H_2_S signaling involves HO/CO-mediated activation of TRPV4 or if TRPV4-mediated Ca^2+^ influx activates HO (Boehning et al., 2004), pressurized mesenteric arteries were exposed to either increasing concentrations of CORM-3 in the presence or absence of the TRPV4 inhibitor GSK219 or to the TRPV4 agonist GSK101 in the presence or absence of CrMP. Administration of CORM-3 elicited concentration-dependent vasodilation and increased endothelial Ca^2+^ events in a TRPV4-dependent manner (Figure 3A, B). However, direct activation of TRPV4 elicited robust vasodilation that was unaffected by inhibition of HO (Figure 3C). These findings place HO signaling functionally upstream of TRPV4 activation.

**Figure 3.**
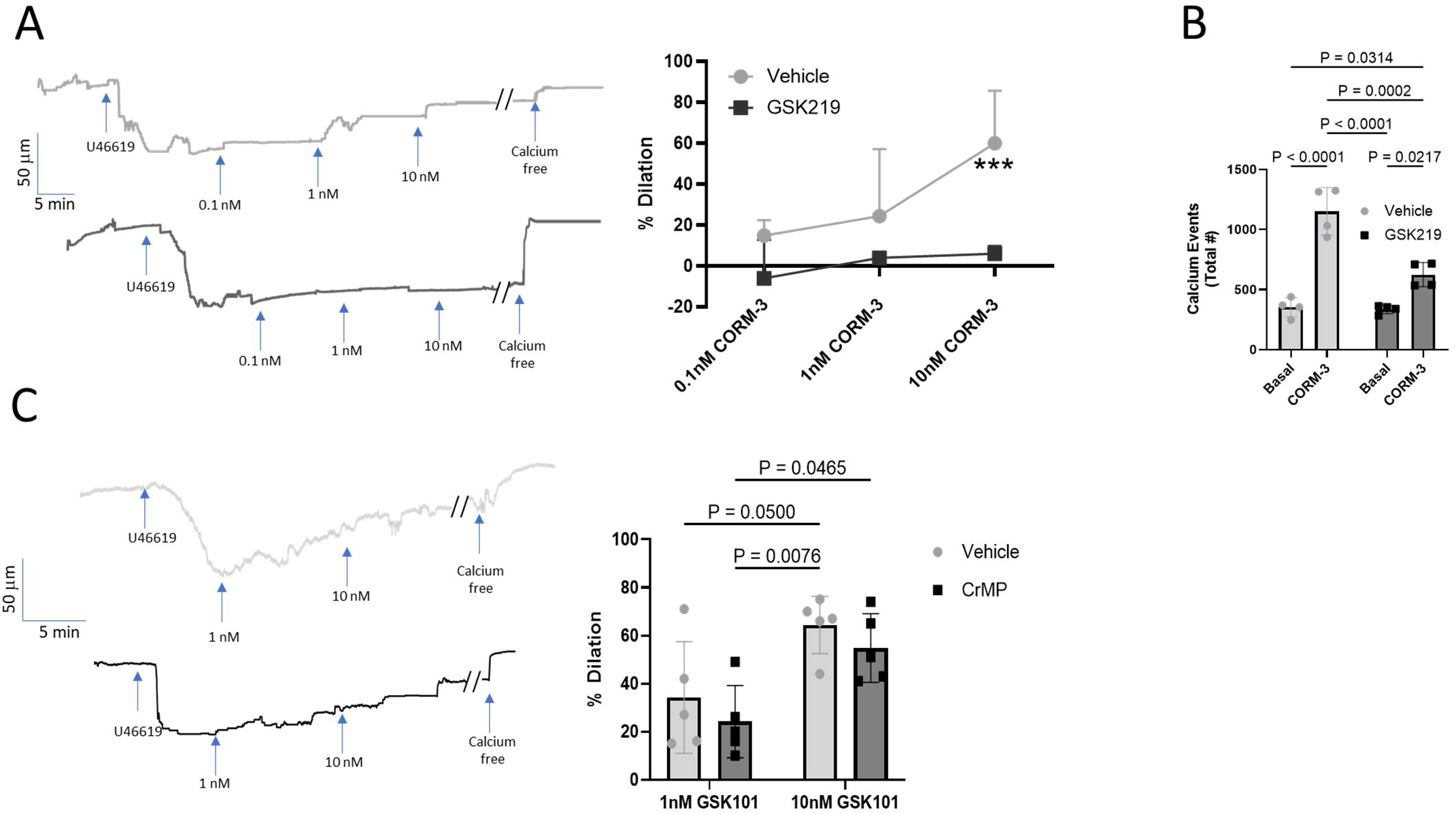
HO-dependent CO signaling activates TRPV4-mediated endothelial Ca²⁺ influx and vasodilation in pressurized rat mesenteric arteries (150 – 200 μm), whereas direct TRPV4 activation elicits vasodilation independently of HO. (**A**) Representative traces and summary data. Arteries were exposed to increasing concentrations of CORM-3 (0.1, 1, and 10 nM) in the presence and absence of the TRPV4 inhibitor GSK219 (300 nM). (**B**) Endothelial cells of pressurized mesenteric arteries were loaded with the fluorescent Ca²⁺ indicator Fluo-4 AM (5 µM), and endothelial Ca²⁺ events were measured under baseline conditions and following exposure to CORM-3 (10 nM), in the presence or absence of GSK219 (300 nM). (**C**) Representative traces and summary data. Isolated arteries were exposed to GSK101 (1 and 10 nM) in the presence and absence of the HO-2 inhibitor CrMP (1 μM). Data are presented as mean ± SD (n = 5/group) and analyzed by 2-way ANOVA. ***P<0.001.

### HO-2 and TRPV4 channels colocalize in endothelial cells

To examine the spatial association of HO-2 and TRPV4, proximity ligation assays (PLA) were performed in human aortic endothelial cells. Representative PLA images demonstrated positive puncta, indicating close spatial association (<40 nm) between HO-2 and TRPV4 channels (Figure 4). Minimal signal was observed in negative control preparations lacking primary antibodies, supporting the specificity of the PLA labeling. These findings suggest that HO-2/TRPV4 reside within closely associated endothelial signaling microdomains.

**Figure 4.**
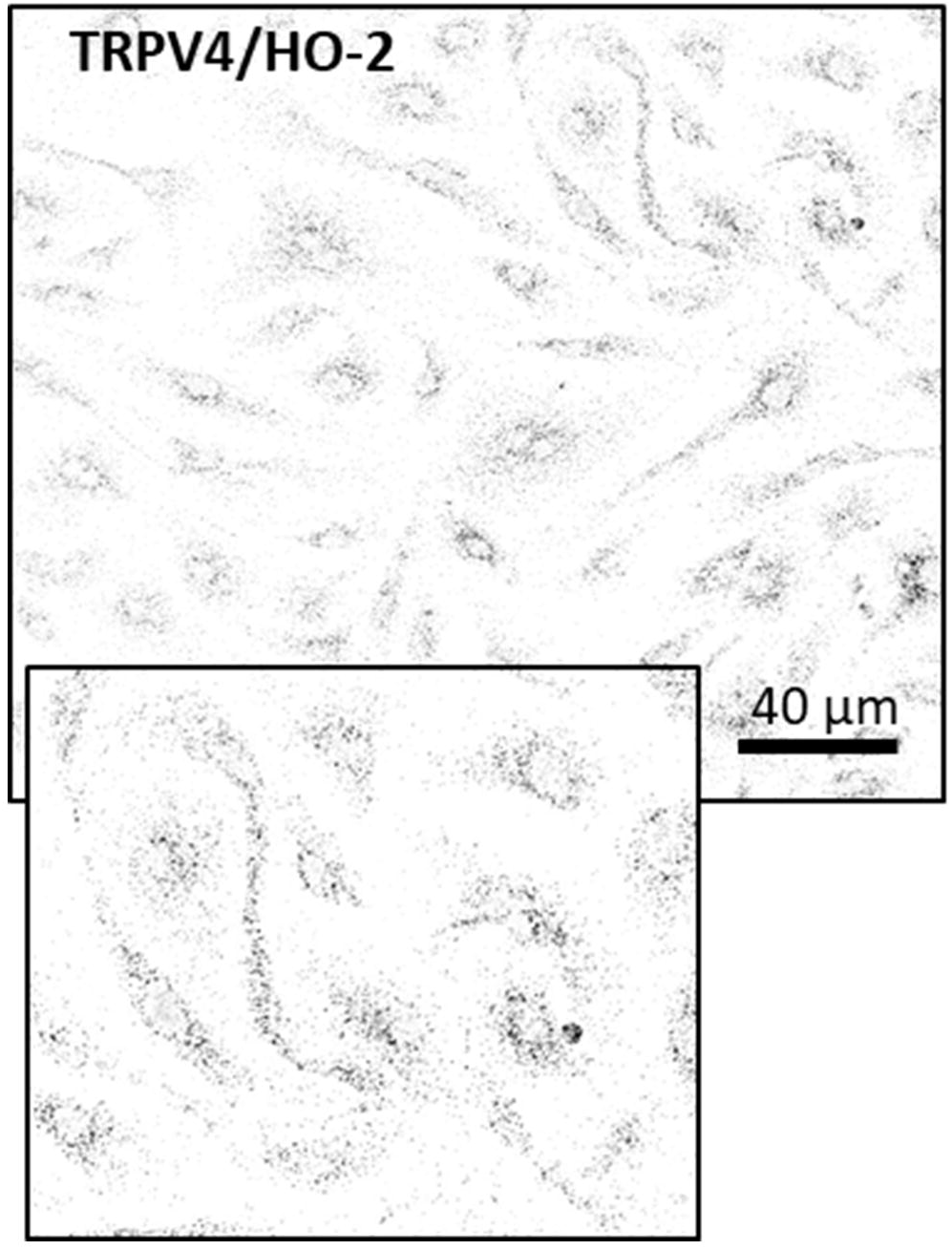
Representative Duolink proximity ligation assay images demonstrating protein–protein interactions between HO-2 and TRPV4 channels in human aortic endothelial cells. PLA-positive signals (black puncta) indicate spatial proximity (<40 nm) between target proteins. Probe-only controls, in which primary antibodies were omitted, exhibited minimal background PLA signal.

### CO-mediated vasodilation is independent of BK channel activation

In the present study, we show that H₂S-induced vasodilation requires HO activation, suggesting that CO acts as a downstream signaling intermediate in this response. As CO has been reported to activate BK channels directly and promote vasodilation (Leffler et al., 2011; R. Wang & Wu, 1997), we examined whether endothelial BK channels contribute to CO-induced dilation. Isolated mesenteric arteries were exposed to increasing concentrations of the CO donor CORM-3 in the presence or absence of luminal IBTx. Luminal inhibition of endothelial BK channels did not significantly alter CORM-3–induced vasodilation (Figure 5), suggesting that endothelial BK channels are unlikely to be a major downstream target of CO in this preparation.

**Figure 5.**
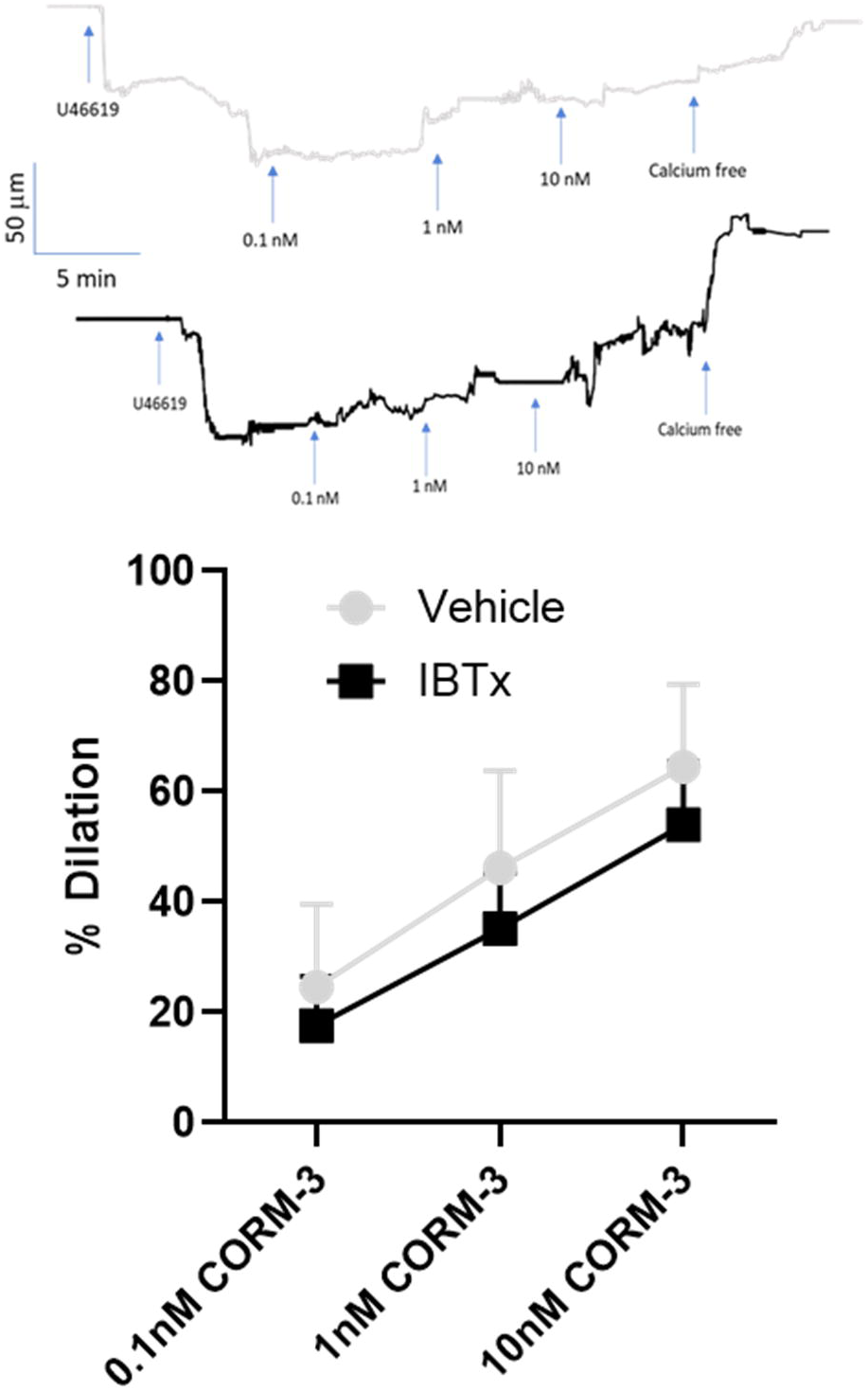
Representative traces and summary data. Inhibition of BK channels does not inhibit CO-induced vasodilation in pressurized rat mesenteric arteries (150 – 200 μm). Arteries were exposed to increasing concentrations of CORM-3 (0.1, 1, and 10 nM) in the presence and absence of BK inhibitor IBTx (Lumen, 100 nM). Data are presented as mean ± SD (n = 5/group) and analyzed by 2-way ANOVA.

### SK/IK channels contribute to CO-mediated vasodilation

We and others have shown that Ca^2+^ influx through TRPV4 channels can activate SK and IK channels but not BK channels (Lin et al., 2015; Naik & Walker, 2018; Ottolini et al., 2020). To determine whether endothelial SK/IK channels contribute to CO-mediated vasodilation, arteries were treated with CORM-3 in the presence or absence of the SK/IK inhibitors TRAM-34 and apamin. Inhibition of SK/IK channels significantly attenuated CO-mediated vasodilation compared with vehicle-treated arteries (Figure 6). These findings indicate that SK/IK channel activity contributes substantially to downstream endothelial signaling following CO stimulation.

**Figure 6.**
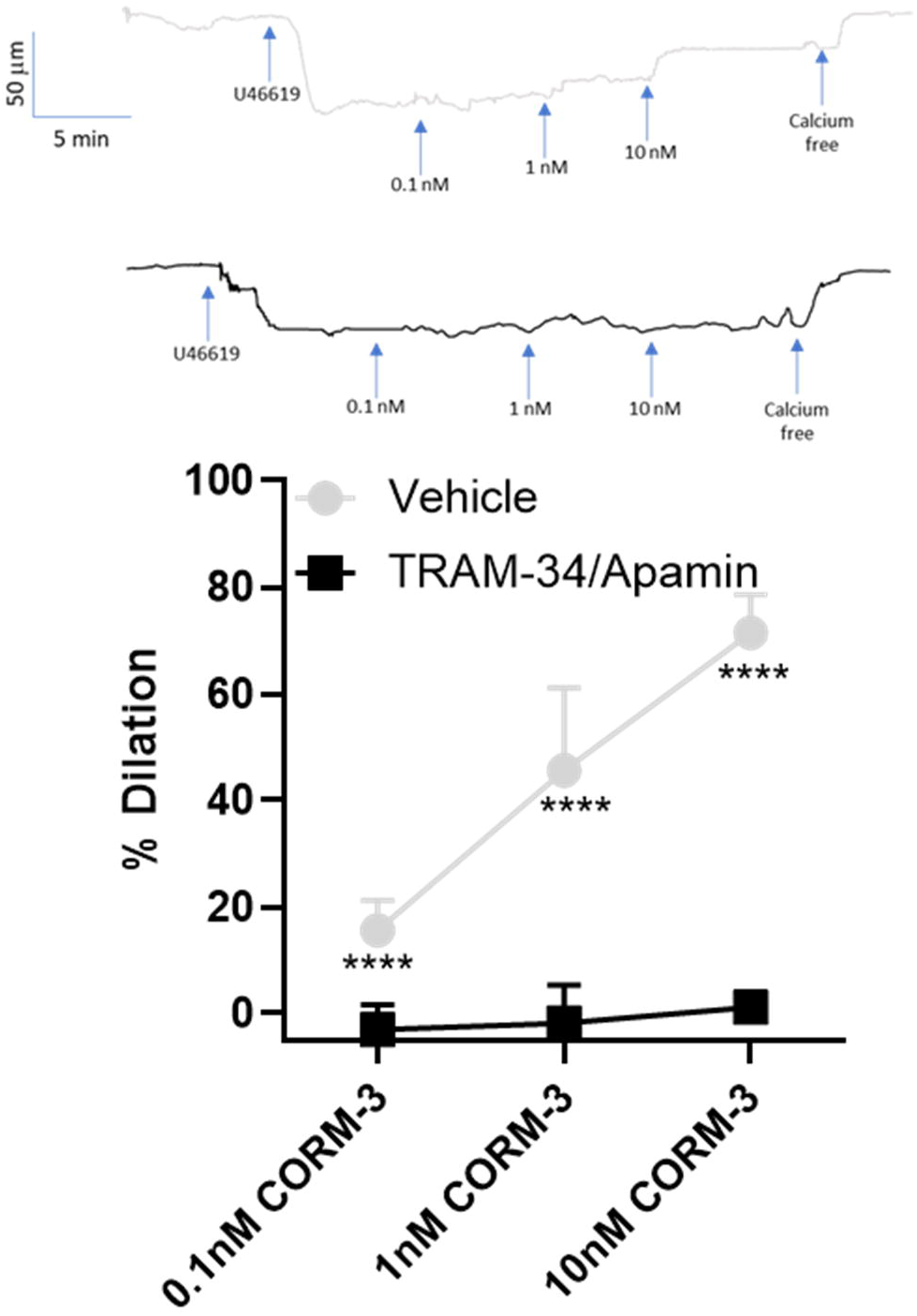
Representative traces and summary data. Inhibition of SK/IK channels abolishes CO-induced vasodilation in pressurized rat mesenteric arteries (150 – 200 μm). Arteries were exposed to increasing concentrations of CORM-3 (0.1, 1, and 10 nM) in the presence and absence of SK/IK inhibitors, TRAM-34 (1 μM)/Apamin (100 nM), respectively. Data are presented as mean ± SD (n = 5/group) and analyzed by 2-way ANOVA. ****P<0.0004 TRAM-34/Apamin vs. Vehicle.

## 4 Discussion

Previous studies have shown that endogenous H_2_S is an endothelial-derived vasodilator and a putative endothelial-derived hyperpolarizing factor (Tang et al., 2013). However, the signaling mechanisms linking H_2_S to TRPV4 activation remain incompletely defined. Our findings demonstrate that heme oxygenase signaling is required for H₂S-mediated vasodilation and support a model in which H₂S stimulates HO-2-derived CO production to engage TRPV4-dependent endothelial signaling. The major findings of the present study are (1) HO inhibition abrogates H_2_S-mediated vasodilation, whereas repletion of CO during HO inhibition restores the vasodilatory response, (2) TRPV4 inhibition abolishes CO-mediated vasodilation, and (3) inhibition of SK/IK but not BK channels prevents CO-mediated vasodilation. Taken together, these findings identify HO-derived CO as a previously unrecognized intermediary linking H_2_S signaling to endothelial TRPV4 activation and vasodilation (Figure 7).

**Figure 7.**
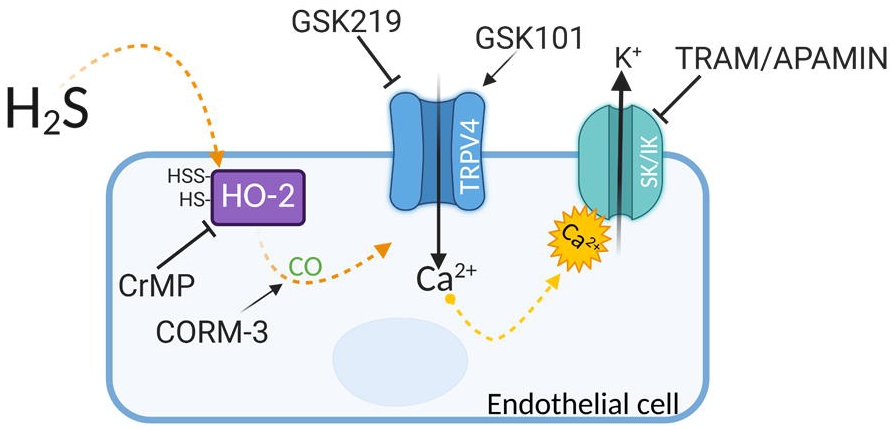
Graphical summary. H_2_S-mediated vasodilation involves sulfhydration and activation of HO-2. The resulting increase in CO production activates TRPV4 channels. TRPV4 sparklets activate SK/IK channels. Created in BioRender. Naik, J. (2026) https://BioRender.com/t2yna8k

HO-derived CO acts as a vital gasotransmitter in the cardiovascular system by regulating vascular tone, suppressing inflammation, and preventing abnormal cell growth. Our findings suggest that CO may serve as a critical signaling molecule in mediating H_2_S-induced vasodilation. Indeed, restoration of vasodilation by exogenous CO in the presence of HO inhibition further supports the concept that CO production is necessary for H_2_S signaling. These findings are consistent with previous reports demonstrating that H_2_S can enhance HO activity by interacting with ferric heme intermediates, thereby increasing CO production (Matsui et al., 2018). Although the two principal isoforms of HO are both expressed in EC (Rydkina et al., 2002), the majority of studies investigating the intersection of the heme oxygenase and H_2_S pathways have identified a relationship between H_2_S and HO-1. Indeed, NaHS administration has been shown to increase HO-1 expression and CO levels (Qingyou et al., 2004; Ren et al., 2026; Zhang et al., 2015). However, multiple lines of investigation suggest that HO-2 is the most likely downstream target of H_2_S. Under basal conditions, HO-2 is the dominant isoform in EC (Bellner et al., 2009; Haider et al., 2002; Krönke et al., 2007; Parfenova et al., 2001). Moreover, the primary mechanism by which H_2_S exerts its physiological effects is through direct protein modification. H_2_S sulfhydrates cysteine residues on target proteins, forming hydroperoxide-sulfide moieties. H_2_S has been shown to sulfhydrate numerous proteins (Mustafa et al., 2011; Paul & Snyder, 2015), including ion channels in the vasculature. For example, H_2_S-induced vasodilation has been linked to sulfhydration of Cys43 on the Kir6.1 subunit of the ATP-dependent K^+^ channel (KATP) in vascular smooth muscle cells (Mustafa et al., 2011). Furthermore, the HO-1 sequences for rats and humans (UniProt IDs: P06762, P09601) do not contain cysteine residues and therefore cannot be activated by H_2_S via persfuldidation. In contrast, rat (UniProt ID: Q07523) and human HO-2 (UniProt: P30519) contain 8 and 4 cysteine residues, respectively.

We previously showed that H_2_S can sulfhydrate TRPV4 channels, increasing TRPV4-mediated Ca^2+^-events and inducing vasodilation (Komagiri, 2024; Vergnolle et al., 2014; Zhou et al., 2023; Naik et al., 2016). In the present study, we show that H_2_S sulfdydrates HO-2 and increases CO production. Therefore, either TRPV4 or HO-2 may serve as the proximal target of H_2_S signaling. Boehning et al. identified three possible 1–10 calmodulin-binding motifs in HO-2 and localized functional binding primarily to the motif around HO-2 residues 66–75. They compared this region with HO-1 and found that only the HO-2 motif had the required sequence and α-helical positioning; mutation of the critical HO-2 residues abolished calmodulin binding and Ca²⁺-dependent activation (Boehning et al., 2004). Therefore, H_2_S may activate HO-2 not through direct sulfhydration but secondary to TRPV4 activation. To test this possibility, we examined vasodilation induced by direct pharmacological activation of TRPV4 in the presence and absence of HO inhibition. We found that HO inhibition did not affect TRPV4-dependent vasodilation. In contrast, vasodilation induced by the CO donor was abolished by TRPV4 channel inhibition. Furthermore, we demonstrate in the present study that TRPV4 and HO-2 colocalize in EC. Taken together, these findings suggest that HO-2 is a direct target of H_2_S, which sulfhydrates the protein, increases CO production, and subsequently activates TRPV4.

HO-2 and TRPV4 channels have been shown to form an essential and functional complex in subfornical organ neurons that mediated hypoxia-induced increases in intracellular calcium (F. Yang et al., 2016), suggesting that HO/CO may be required for TRPV4 activation. Our observation that HO inhibition abolished H_2_S-mediated vasodilation but had no effect on direct TRPV4 activation suggests that HO signaling acts upstream of TRPV4. However, the molecular mechanism by which CO activates TRPV4 remains unknown. CO may indirectly stimulate TRPV4 through changes in arachidonic-acid metabolism and EET formation, redox-sensitive signaling, or kinase-dependent channel activation.

11,12-EET has been shown to activate vascular TRPV4 channels (Earley et al., 2005). In rats, 11,12-EETs are produced by cytochrome (CYP) P450 2C11 (Liu et al., 2019). CO may increase 11,12-EET production; however, this appears unlikely, as during the normal CYP catalytic cycle, the heme iron is reduced from Fe³⁺ to Fe²⁺, after which molecular oxygen binds to the ferrous heme. CO competes with oxygen for this Fe²⁺ heme iron and forms a stable CYP–Fe²⁺–CO complex, preventing oxygen binding and activation and interrupting substrate oxidation (Munro et al., 2018).

Physiological concentrations of CO generate mitochondrial reactive oxygen species (ROS) (Bilban et al., 2006; Zuckerbraun et al., 2007). Previous studies have shown that ROS can increase intracellular Ca^2+^ in endothelial cells (Gandhirajan et al., 2013). Indeed, H_2_O_2_ has been shown to increase TRPV4-dependent Ca^2+^-influx in pulmonary artery endothelial cells (Suresh et al., 2015).

CO may activate intracellular kinase pathways through soluble guanylyl cyclase (Naik & Walker, 2003). Fan et. al. demonstrated that TRPV4 channels are activated by serine phosphorylation, with the protein kinase C activator phorbol 12-myristate 13-acetate (Fan et al., 2009). cGMP can activate PKA by inhibiting cGMP-dependent phosphodiesterase (Zaccolo & Movsesian, 2007). Furthermore, PKA-dependent phosphorylation at S824 increased TRPV4 Ca^2+^-influx (Cao et al., 2018). Although these mechanisms are of considerable interest, their elucidation will require further investigation.

K⁺ channel activation is a critical component of H₂S-induced vasodilation. Consistent with this concept, elevation of extracellular K⁺ abolishes H₂S-induced vasodilation, and sulfhydration of smooth muscle KATP channels and endothelial IK channels has been implicated in the vasodilatory response to H₂S (Cheng et al., 2004; Mustafa et al., 2011). We previously demonstrated that H₂S-induced dilation and endothelial Ca²⁺ events in rat mesenteric arteries are blocked by TRPV4 inhibition (Naik et al., 2016), indicating that TRPV4 channels are an important component of H₂S signaling. We also found that vasodilation induced by either H₂S or direct pharmacological activation of TRPV4 channels with GSK101 was abolished by endothelial BK-channel inhibition (Jackson-Weaver et al., 2011; Naik et al., 2016). In contrast, the present study demonstrates that dilation to a CO donor, a downstream mediator of H₂S signaling, is abolished by SK/IK channel inhibition but unaffected by BK channel inhibition. These findings indicate that the specific K⁺ channels recruited downstream of TRPV4 activation may depend on the mode, magnitude, or subcellular localization of TRPV4-mediated Ca²⁺ influx.

Direct pharmacological activation of TRPV4 channels with GSK101 may generate a larger or more spatially extensive Ca²⁺ influx than physiological activation by CO, thereby reaching the Ca²⁺ concentrations required to activate endothelial BK channels. In contrast, SK and IK channels, which possess higher intrinsic Ca²⁺ sensitivity through constitutively associated calmodulin, may be preferentially activated by the smaller or more localized Ca²⁺ signals generated by CO-dependent TRPV4 activation. Indeed, SK and IK channels are high-affinity, voltage-independent Ca²⁺ sensors with half-maximal activation at approximately 300–500 nM intracellular Ca²⁺, whereas BK-channel activation generally requires micromolar local Ca²⁺ concentrations and is strongly influenced by membrane potential and auxiliary β subunits (Ledoux et al., 2006; Xia et al., 1998). The Ca²⁺ and voltage sensitivity of endothelial BK channels remains to be characterized, and their auxiliary subunit composition is currently unknown.

Consistent with the ability of TRPV4 channels to couple to multiple K⁺-channel pathways, we previously showed that GSK101-induced dilation in gracilis arteries is inhibited by SK/IK-channel blockade (Naik & Walker, 2018), and Sonkusare et al. demonstrated that GSK101-induced vasodilation in mesenteric arteries is abolished by SK/IK inhibition (Sonkusare et al., 2012). Conversely, Earley et al. showed that 11,12-EET activates TRPV4 channels and produces smooth muscle BK channel-dependent dilation in cerebral arteries (Earley et al., 2005). Taken together, these observations suggest that TRPV4 channels do not couple exclusively to a single endothelial K⁺-channel subtype. Instead, coupling to BK versus SK/IK channels likely varies with vascular bed, activating stimulus, Ca²⁺-signal amplitude, and the spatial organization of TRPV4 channels with downstream effectors. In the present study, the lack of an effect of BK-channel inhibition and the abolition of CO-induced dilation by SK/IK blockade support a signaling pathway in which H₂S stimulates HO-dependent CO production, CO activates endothelial TRPV4 channels, and the resulting Ca²⁺ influx preferentially activates SK/IK channels to produce membrane hyperpolarization and vasodilation.

To our knowledge, this is the first study to demonstrate a functional association between HO-2 and SK/IK channels. In comparison, others have linked HO-1 to Ca^2+^-activated K^+^ channels in other cell types. For example, in glomus cells and HEK293 cell expression systems, the Ca^2+^-dependent HO-2 isoform has been shown to associate with and be activated by BK channels (Boehning et al., 2004; Williams et al., 2004). In addition, hemin-induced HO-1 upregulation potentiates EDH-type relaxations in the mesenteric artery of the SHR. The improvement in EDH-type responses induced by HO-1 can be attributed to increased expression of Na^+^-K^+^-ATPase, facilitating relaxation via the IK_Ca_-Na^+^-K^+^-ATPase pathway. This beneficial effect may be mediated by HO-1’s antioxidant properties, as suggested by bilirubin production (Li et al., 2013).

In conclusion, the present findings support a model in which H_2_S stimulates HO-2-derived CO production that subsequently activates TRPV4-dependent endothelial signaling to promote vasodilation in resistance-sized mesenteric arteries. These findings identify HO-derived CO as a critical intermediary in H_2_S signaling pathways.

## Conflict of Interest

The authors declare that the research was conducted in the absence of any commercial or financial relationships that could be construed as a potential conflict of interest.

## Author Contributions

J.S.N. conceived and designed research; J.R.A. and C.X.N performed all experiments; J.R.A., C.X.N and J.S.N analyzed data; J.R.A., C.X.N., L.V.G.B., and J.S.N. interpreted results of experiments; J.R.A., C.X.N., and J.S.N. prepared figures; J.R.A. wrote first draft of the manuscript; J.R.A., C.X.N., L.V.G.B., and J.S.N. edited and revised manuscript; J.R.A., C.X.N., L.V.G.B., and J.S.N. approved final version of manuscript.

## Funding

R01 HL160606-04 (J.S.N), UNM Comprehensive Cancer Center Advanced Light Microscopy Shared Resource, NCI 2P30 CA118100.

## Data Availability Statement

The data that support the findings of this study are available from the corresponding author upon reasonable request.

## Notes

### Competing Interest Statement

The authors have declared no competing interest.

