## Supplementary figures and images for "Hydrogen sulfide-mediated vasodilation requires heme oxygenase-derived carbon monoxide"

### Supplemental Figure 1

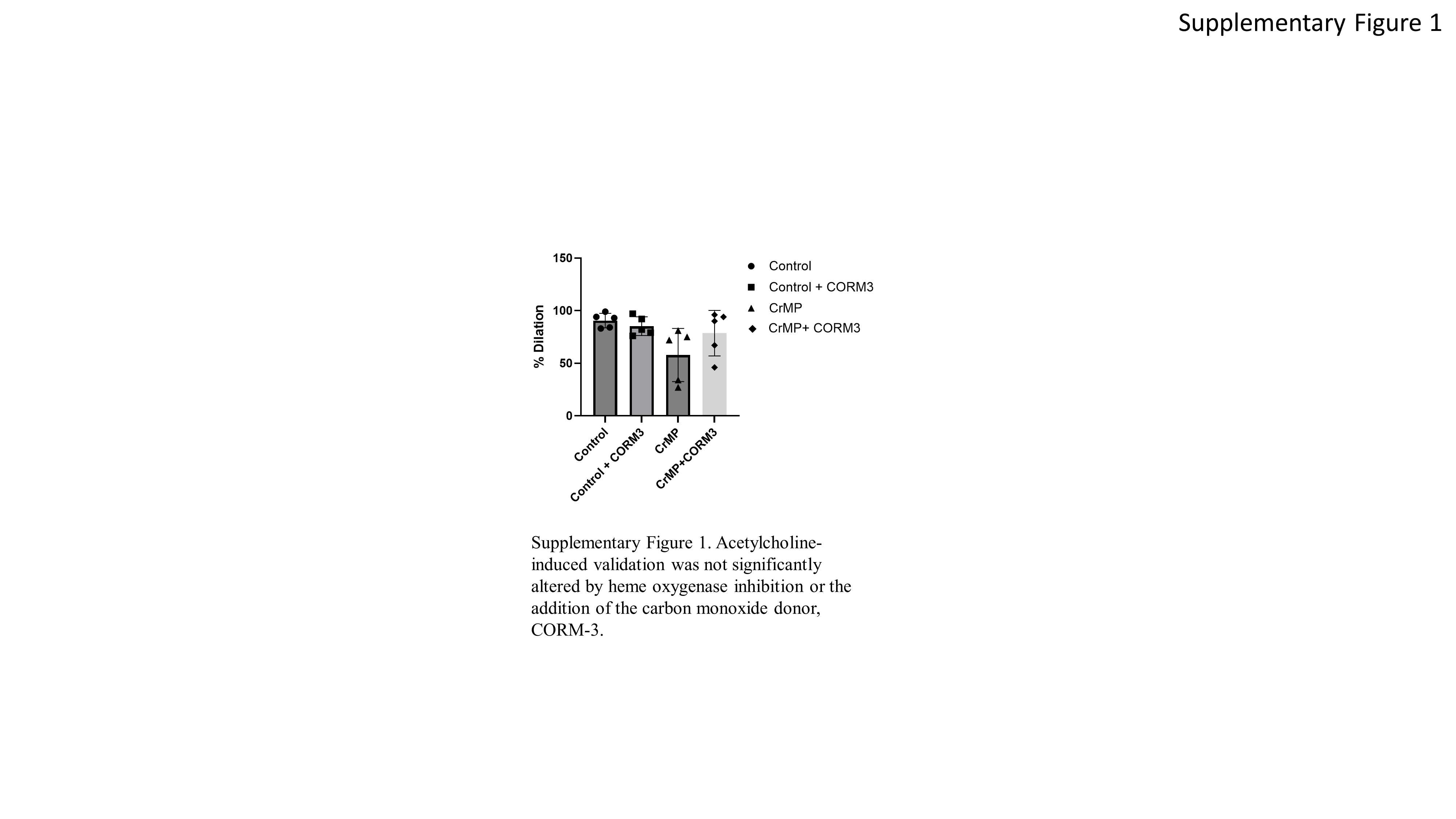
